# Specific effect of a dopamine partial agonist on counterfactual learning: evidence from Gilles de la Tourette syndrome

**DOI:** 10.1101/064345

**Authors:** Alexandre Salvador, Yulia Worbe, Cécile Delorme, Giorgio Coricelli, Raphaël Gaillard, Trevor W. Robbins, Andreas Hartmann, Stefano Palminteri

**Author notes:** **Corresponding authors:** Stefano Palminteri, PhD. **Author contributions** AS performed the experiments, analysed the data, interpreted the results and wrote the manuscript. YW designed the study, performed the experiment, interpreted the results and edited the manuscript. CD performed the experiments and edited the manuscript. GC, RG, TWR interpreted the results and edited the manuscript. AH designed the study and interpreted the results. SP designed the study, analysed the data, interpreted the results and wrote the manuscript.

## Abstract

The dopamine partial agonist aripiprazole is increasingly used to treat pathologies for which other antipsychotics are indicated because it displays fewer side effects, such as sedation and depression-like symptoms, than other dopamine receptor antagonists. Previously, we showed that aripiprazole may protect motivational function by preserving reinforcement-related signals used to sustain reward-maximization behaviour in a simple action-outcome learning task. However, the effect of aripiprazole on more cognitive facets of human reinforcement learning, such as learning from the hypothetical outcomes of alternative courses of action (i.e., counterfactual learning), is unknown.

To test the influence of aripiprazole on counterfactual learning, we administered a reinforcement-learning task that involves both direct learning from obtained outcomes and indirect learning from forgone outcomes to two groups of Gilles de la Tourette (GTS) patients, one consisting of patients who were completely unmedicated and the other consisting of patients who were receiving aripiprazole monotherapy, and to healthy subjects. We replicated a previous finding showing that aripiprazole does not affect direct learning from obtained outcomes in GTS. We also found that whereas learning performance improved in the presence of counterfactual feedback in both healthy controls and unmedicated GTS patients, this was not the case in aripiprazole-medicated GTS patients.

Our results suggest that whereas aripiprazole preserves direct learning of action-outcome associations, it may impair more complex inferential processes, such as counterfactual learning, from forgone outcomes.

## Introduction

Aripiprazole is a recently introduced antipsychotic medication with dopamine receptor partial agonist mechanisms^1^. In schizophrenia patients, aripiprazole exhibited efficacy comparable to that of typical and atypical antipsychotics in the treatment of positive and negative symptoms as well as in the prevention of relapse^2,3^. Its efficacy has also been demonstrated in other neurological disorders for which antipsychotics are indicated, such as Gilles de la Tourette syndrome (GTS)^4,5^. Its tolerability is often considered superior to that of typical antipsychotics, and it is associated with fewer adverse side effects, such as extrapyramidal and metabolic syndromes^6,7^. In GTS, aripiprazole is effective for suppressing tics while displaying a less severe side effect profile than dopamine receptor antagonists with regard to motivational deficits such as sedation and depressive reactions^8^. Due to this advantageous cost-benefit trade-off, aripiprazole has become a widely prescribed treatment for schizophrenia and GTS.

Pharmacological studies in humans suggest that dopamine receptor antagonist-induced sedation and depressive states may be the consequence of blunting of reward-related signals^9,10^. For instance, reward-seeking behaviour and reward prediction errors encoded in the ventral striatum were reduced by haloperidol administration in healthy volunteers^11^. In contrast, both functions were preserved in GTS patients medicated with aripiprazole^12^. These results are consistent with the idea that, as opposed to dopamine receptor antagonists, aripiprazole preserves motivational functions via preserving reward-related signals.

Despite the observations described above, in humans, reinforcement learning is seldom solely based on learning from obtained outcomes (i.e., factual outcomes)^13^. In fact, human reinforcement learning often makes use of more abstract inferential processes, including learning from counterfactual outcomes^14,15^. In other words, counterfactual learning refers to the ability to learn not only from direct experience but also from hypothetical outcomes (the outcomes of the option that was not chosen). Previous neuroimaging studies have demonstrated that subjects take into account counterfactual feedback when it is available and have suggested that this form of learning could be underpinned by a dorsal prefrontal system^16-18^, despite the fact that some other studies have suggested that factual and counterfactual outcomes may be processed by the same neural system involving the ventral striatum^19,20^. On the pharmacological level, a recent study that investigated counterfactual learning under a regimen of dietary depletion of serotonin and dopamine/noradrenalin precursors indicated that brain tracking of fictive prediction error might be sensitive to both dopamine and serotonin manipulation^21^.

Here, to characterize the effect of aripiprazole on counterfactual learning, we administered a recently developed behavioural task that contrasts learning from obtained and hypothetical outcomes to a group of medication-free GTS patients, a group of GTS patients with aripiprazole as their only medication, and matched healthy controls^22^. GTS patients represented an ideal model for this investigation because in this population we have already shown the beneficial effects of aripiprazole compared to a dopamine receptor antagonist in factual learning, that is, in learning from obtained outcomes (as opposed to counterfactual learning, which is learning from forgone outcomes)^12^. In addition, unlike many schizophrenic patients, GTS patients have relatively preserved cognitive efficiency^23,24^. Finally, by investigating the effect of aripiprazole in a population that is commonly treated with this medication, our findings have direct clinical relevance.

## Results

### Demographic and psychometric results

Forty consecutive patients consulting at the GTS Reference Center in Paris were included in the study. Patients taking any medication other than aripiprazole (i.e., pimozide, risperidone, haloperidol, bromazepam or escitalopram) were not included in the final analysis. One patient did not complete the task and was also excluded from the analysis. This led to a final sample of 31 GTS patients (17 without any medication and 14 with aripiprazole monotherapy). In addition, 20 healthy controls obtained through advertising were included. Two significant differences were found when comparing the 3 groups: the number of years of study (F(2,48)=3.2, p=0.050) (although the results of the National Adult Reading Test (NART; F(2,46)=1.2, p=0.31), which measures verbal intelligence, did not differ among the groups) and the Beck Depression Inventory score (BDI, F(2,48)=6.72, p=0.0027). In addition, two scales showed marginal differences among the three groups: the Barratt Impulsiveness Scale (BIS-11; F(2,48)=3.00, p=0.059) and the Behavioural Inhibition / Behavioural Activation Scale (BIS/BAS; F(2,48)=3.04, p=0.057). Flowever, there was no significant difference between the aripiprazole and unmedicated GTS groups on any demographic or psychometric scale or in disease severity index or measures of intelligence (all p>0.2, two-sample t-test).

### Behavioural results

Subjects performed a probabilistic instrumental learning task adapted from previous imaging studies^22,25^. During the instrumental probabilistic learning task, the subjects were required to learn which stimuli had the greatest likelihood of resulting in an advantageous outcome through trial and error. The dependent variable was the subject’s accuracy; an “accurate” (or “correct”) choice was choice of the stimulus that is on average more rewarding or less punishing. Both feedback valence (reward vs. punishment) and feedback information (partial feedback vs. complete feedback) were manipulated using a within subjects factorial design (Figure 1A).

**Figure 1:**
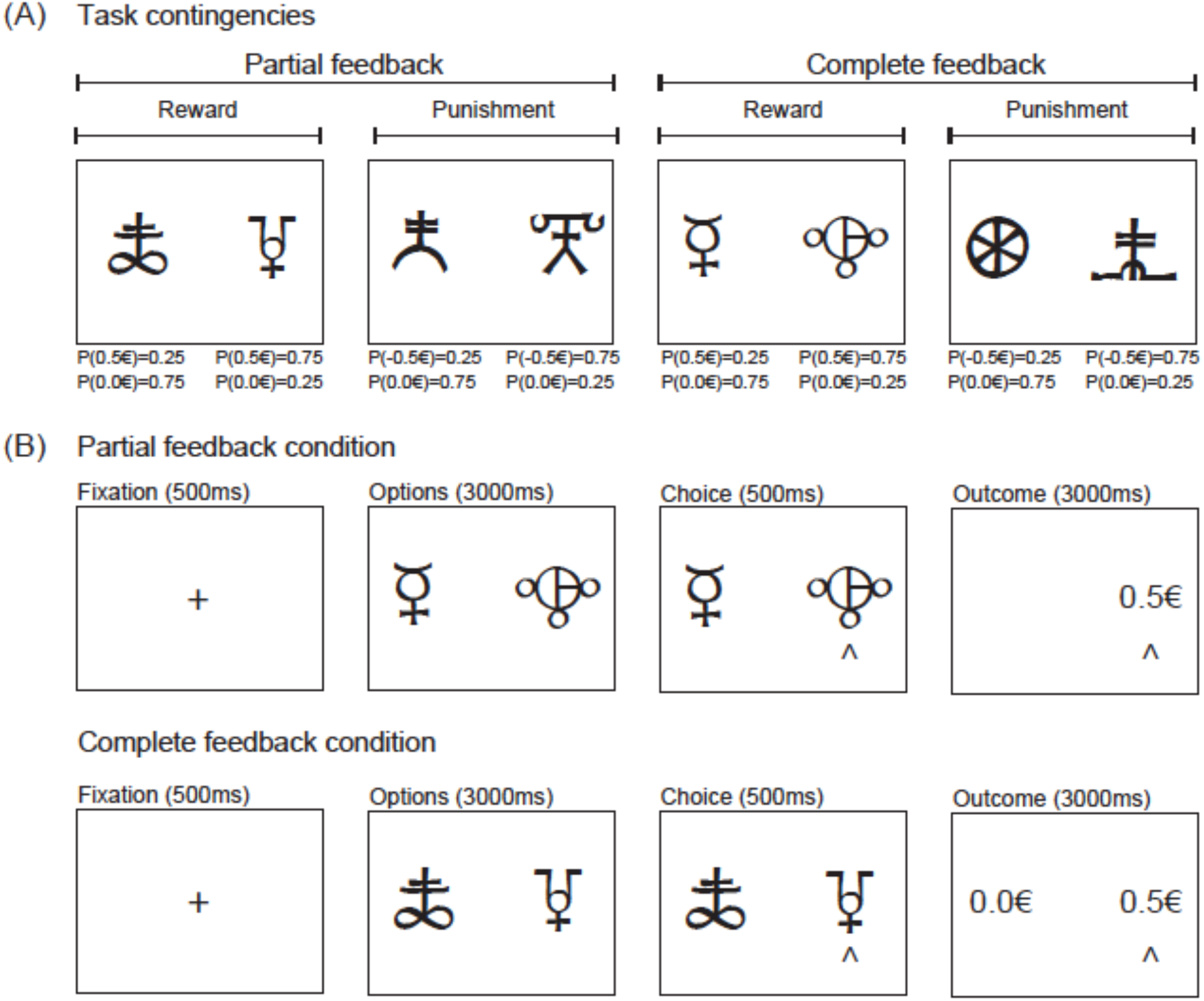
**Task. (A)** Task contingencies and factorial design. **The attribution of symbols to each condition was randomized across subjects**. **(B)** Typical trials. A novel trial started after a fixation screen (500 ms). Subjects were required to select between the two options by pressing one of the corresponding two buttons with their left or right index fingers to select the leftmost or the rightmost option, respectively, within a 3000 ms time window. After the choice window, a red pointer appeared below the selected option for 500 ms. At the end of the trial, the options disappeared and the selected option was replaced by the outcome (“+0.5 €”, “0.0 €” or “-0.5 €”) for 3000 ms. In the complete information context, the outcome corresponding to the unchosen option (counterfactual) was also displayed.

A preliminary analysis of variance (ANOVA) evaluating accuracy as a function of trial, feedback information, feedback valence and group (aripiprazole-treated GTS patients, unmedicated GTS patients, and controls) showed that there was a significant main effect of group (F(2,4864)=14.5, p<0.001), a significant main effect of trial number (F(1,4864)=46.4, p<0.001), and a significant main effect of feedback information (F(1,4864)=23.7, p<0.001), with an interaction effect of group with feedback information (F(2,4864)=15.35, p<0.001) but no main effect of feedback valence (F(1,4864)=0.8, p>0.3). Feedback valence was therefore excluded from the subsequent ANOVA analysis. The final ANOVA, which evaluated accuracy as a function of trial, feedback information, and group, showed that there was a significant main effect of trial number on correct choice rate (F(1,2432)=41.1, p<0.001), indicating that accuracy improved with learning (Figure 2A). In addition, there was a significant main effect of feedback information (F(1,2432)=21.0, p<0.001) and of group (F(2,2432)=12.9, p<0.001), with a significant interaction between group and feedback information (F(2,2432)=13.6, p<0.001), indicating that counterfactual feedback did not positively modulate the accuracy similarly in the three groups. The interaction effect of group and feedback information remained significant when years of study, BDI score, and BIS.11 or BIS.BAS impulsivity scores were included either individually or all together (F(2,2260)=22.8, p<0.001).

**Figure 2:**
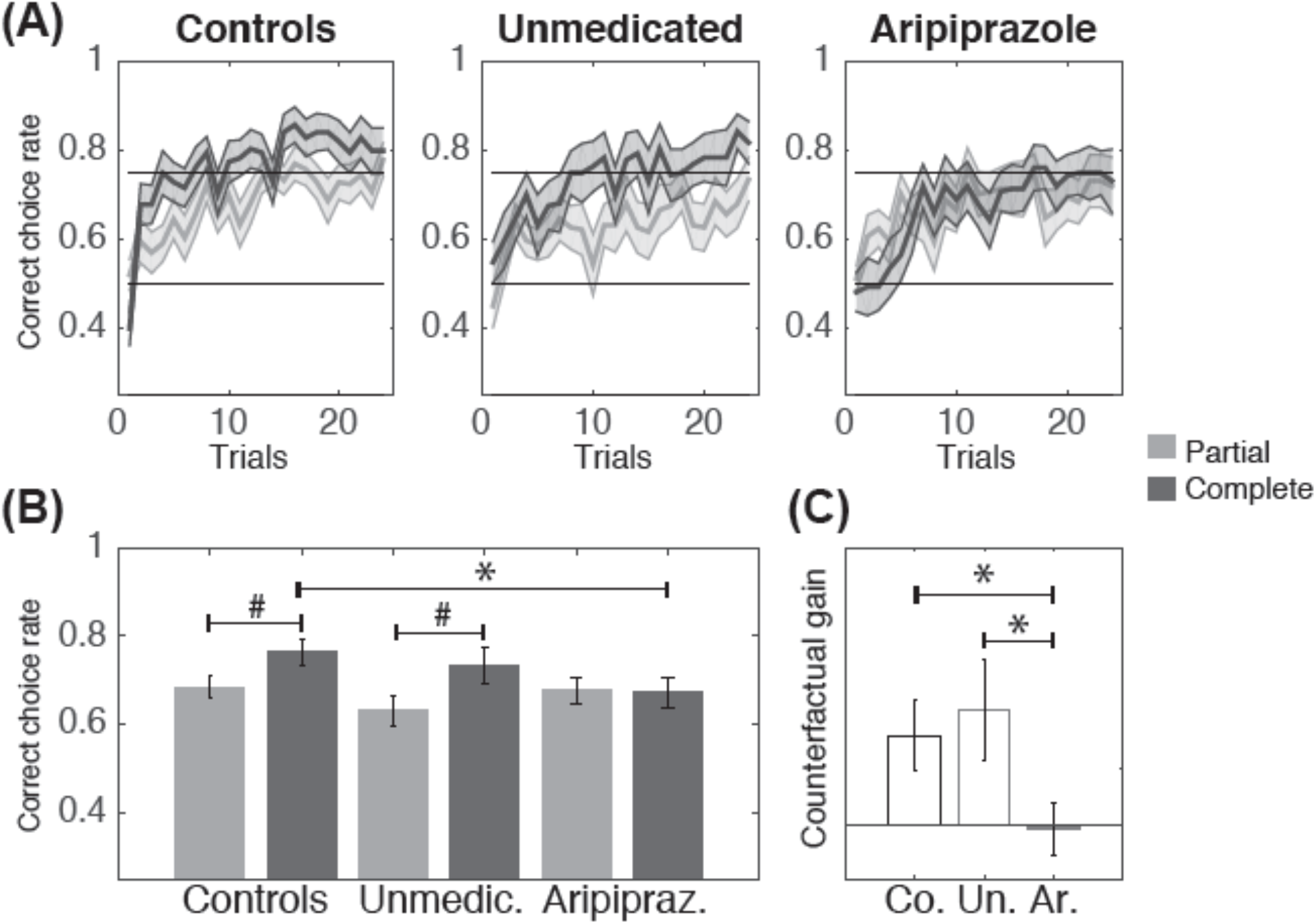
**(A)** Learning curves as a function of feedback information and group. The bold lines represent trial-by-trial average accuracy over the three sessions. The shaded areas between the thin lines represent the standard error of the mean with subject as the random factor. Light and dark grey represent partial and complete feedback conditions, respectively. **(B)** Average performance averaged across the 24 trials. **(C)** Average « counterfactual gain » (complete minus partial accuracy) (Co. = controls; Un. = unmedicated; Ar. = aripiprazole). #p<0.05, one-sample t-test; * p<0.05, two-sample t-test. The error bars represent the standard error of the mean with subject as the random factor.

We then defined the « counterfactual gain » as the difference in accuracy between the complete feedback and the partial feedback conditions. A confirmatory ANOVA evaluating counterfactual gain as a function of group and trial showed a significant main effect of group (F(2,1216)=17.9, p<0.001), indicating that the groups benefited differently from counterfactual information. There was no main effect of trial (F(1,1216)=1.00, p=0.3), indicating that the benefit from counterfactual information was present for the entire duration of the experiment. The effect of group remained significant when years of study, BDI score, and BIS.11 or BIS.BAS impulsivity scores were included either individually or all together (F(2, 1130)=25.1, p<0.001).

To assess the directionality of the critical ANOVA effects in a concise manner, we plotted and analysed the accuracy and the counterfactual gain averaged across trials. Accuracy in the complete feedback condition was significantly higher than accuracy in the partial feedback condition in both the control and unmedicated GTS groups (t(19)=−2.6, p<0.05, and t(16)=−2.3, p<0.05, respectively) but not in the aripiprazole GTS group (t(13)=0.151, p=0.882) (Figure 2B). The average counterfactual gain was significantly higher in the control group and in the unmedicated group than in the aripiprazole group (8.3% controls versus 0.3% aripiprazole, t(31.5)=2.2, p<0.05; t-test 10.5% unmedicated versus 0.3% aripiprazole, t(23.3)=2.1, p<0.05). There was no difference in counterfactual gain between the control and unmedicated groups (t(29.3)=-0.4, p=0.701) (Figure 2C).

Interestingly, in the aripiprazole GTS group, we found a significant correlation between treatment dose and accuracy in the complete feedback condition (Spearman's correlation test p=-0.66, p<0.01), indicating that the higher the dose of atypical antipsychotic medication received, the lower the accuracy in the complete feedback condition. This correlation remained significant when the only patient with a 30-mg dose of aripiprazole was excluded (p=-0.57, p<0.05). In contrast, treatment dose did not correlate with accuracy in the partial feedback condition (Spearman's correlation test p=-0.47, p>0.05). To assess evidence in favour of the null hypothesis (i.e., the absence of counterfactual learning in the aripiprazole group), we performed a Bayesian equivalent of a t-test (see the Materials and Methods section). This analysis showed that the probability that the average counterfactual gain falls within a region of practical equivalence (i.e., ROPE: [−1.5%, 1.5%] interval) is small (4%) in both the unmedicated GTS group and in the control group, whereas this probability is 5.75 times larger (23%) in the aripiprazole group (Figure 4). This Bayesian approach provides evidence that the absence of counterfactual learning in the aripiprazole group does not arise from a lack of power in detecting it.

**Figure 3:**
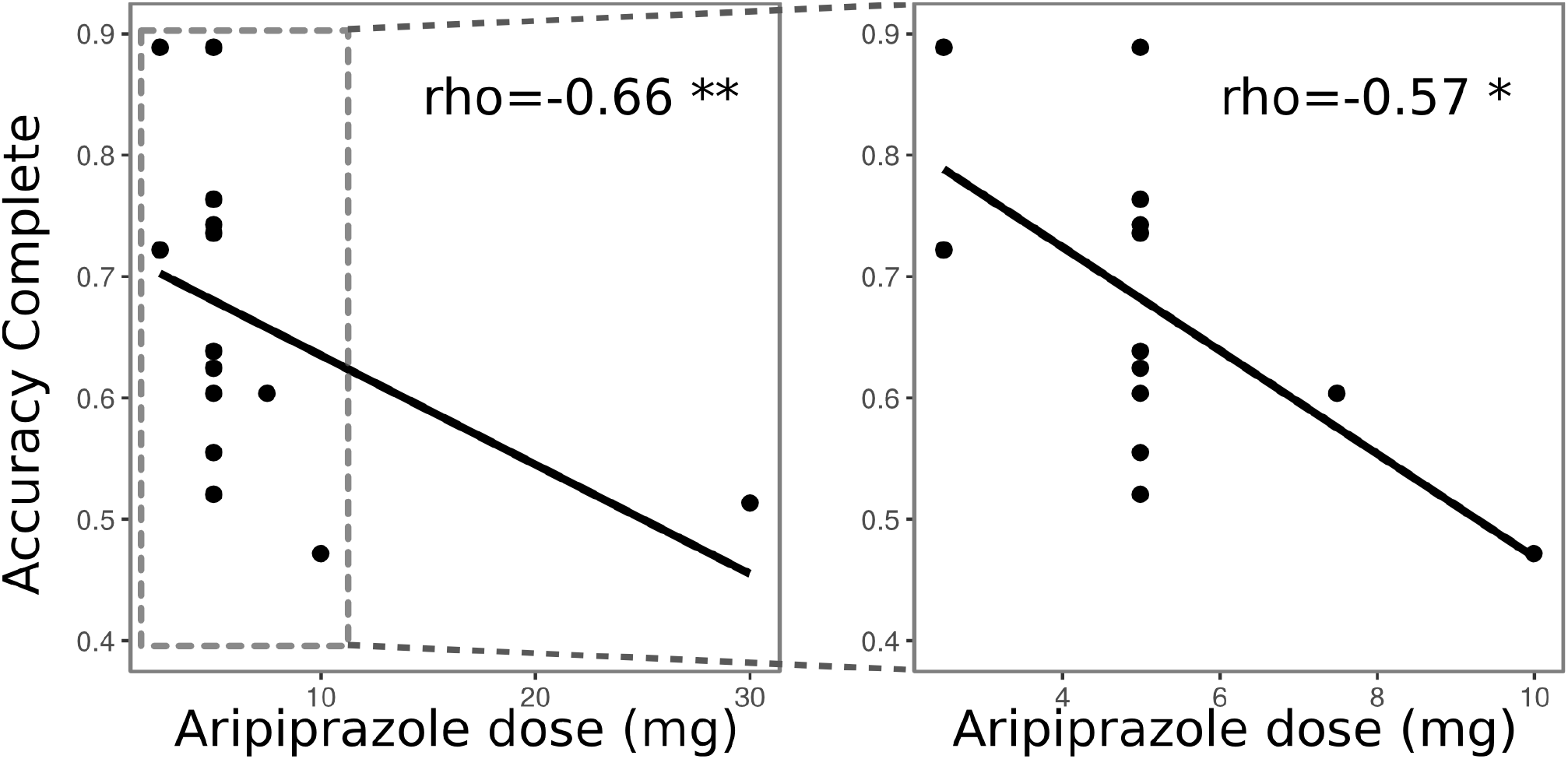
Correlation between aripiprazole dose and accuracy in the complete feedback context in all subjects (left) and excluding the subject with the highest dose (right).

**Figure 4:**
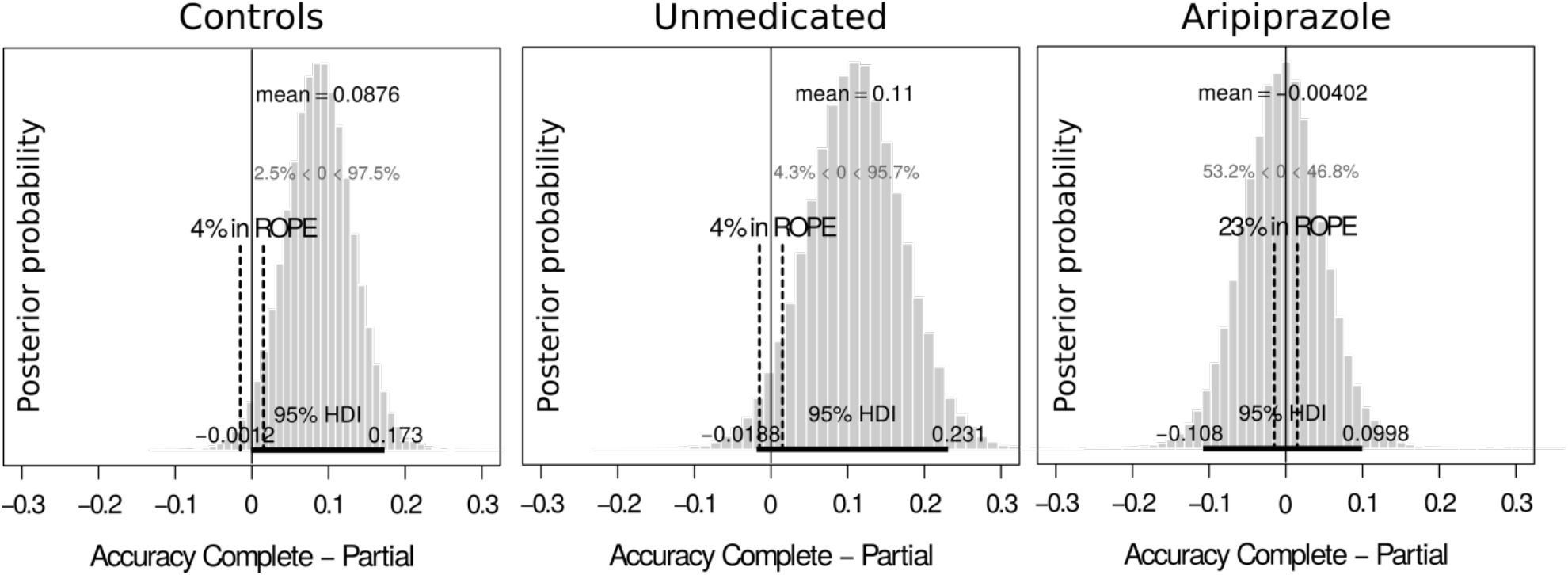
Results of the Bayesian analysis for evidence in favour of the null hypothesis for the difference between accuracy in the complete and partial feedback conditions (i.e., « counterfactual gain ») for each group. ROPE is the region of practical equivalence, which was set at the level of 3% ([−1.5%, 1.5%]. “4% in ROPE” means that there is a 4% chance that the true average counterfactual gain lies within the region of the practical equivalence interval.

Finally, we analysed the reaction times using an ANOVA with trial, group and feedback information as factors (see Table 2). The ANOVA revealed a significant effect of trial, indicating that reaction time decreased across trials (F(1,2432)=17.8, p<0.001). Feedback information and group did not affect reaction time (F(1,2432)=0.08, p=0.772 and F(2,2432)=2.74, p=0.064), nor did any of the interaction factors, indicating that reaction time was not modulated by the pathology or by the treatment.

**Table 1:**
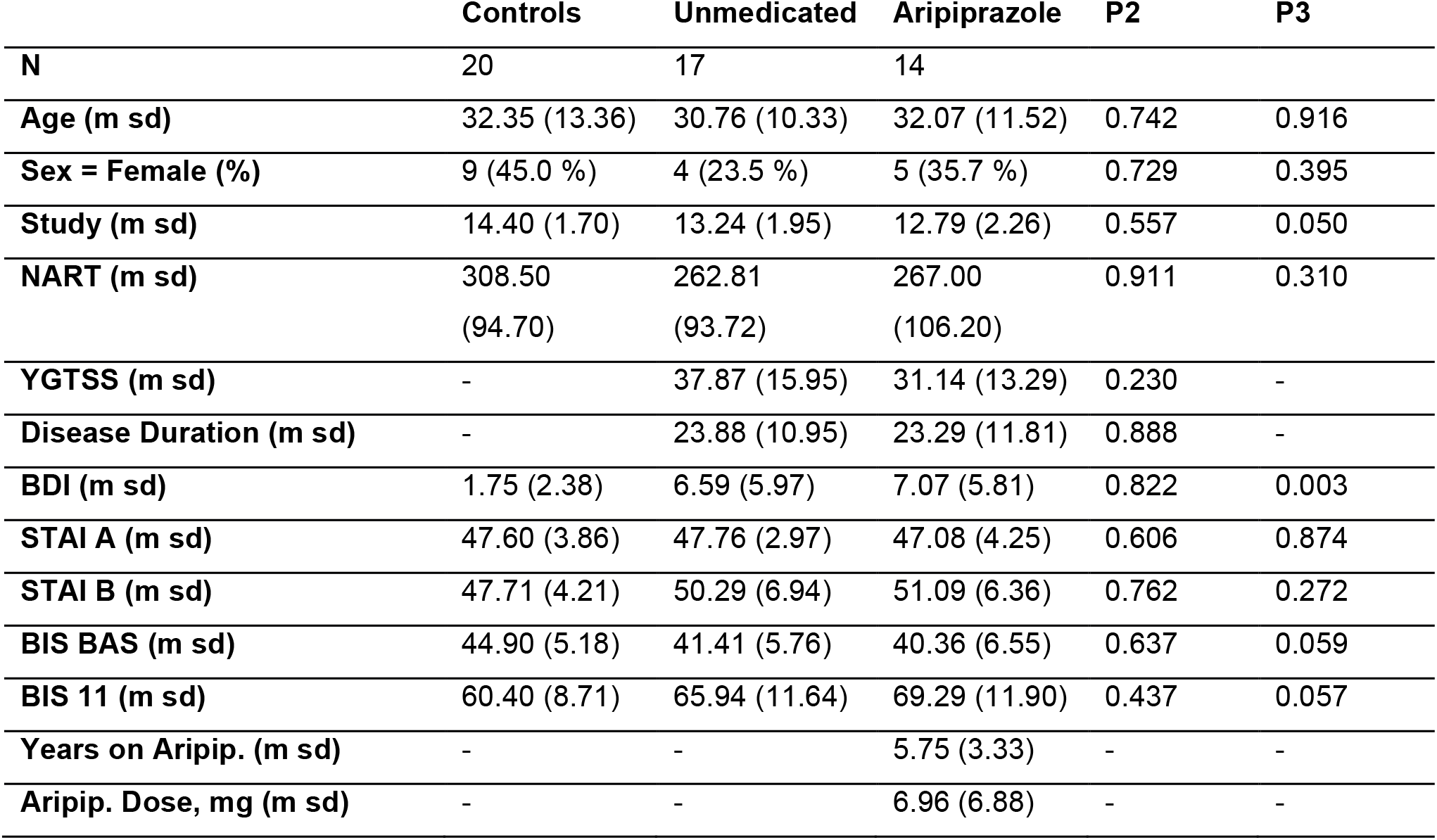
Demographic characteristics and psychometric scales for participants by treatment group. The “p2” p-values correspond to the comparison between the unmedicated and aripiprazole groups (two-sample t-test). The “p3” p-values correspond to the comparison among all three groups (one-way analysis of variance).

**Table 2:** Average behavioural data (mean±s.e.m.).

| Condition | Accuracy (correct choice rate) |  |  |  | Reaction time (seconds) |  |  |  |
| --- | --- | --- | --- | --- | --- | --- | --- | --- |
|  | Rew./Par. | Rew./Com. | Pun./Par. | Pun./Com. | Rew./Par. | Rew./Com. | Pun./Par. | Pun./Com. |
| **Controls** | 0.68±0.04 | 0.76±0.04 | 0.68±0.03 | 0.77±0.04 | 1.16±0.04 | 1.21±0.06 | 1.37±0.06 | 1.27±0.06 |
| **Unmedic.** | 0.61±0.05 | 0.74±0.05 | 0.65±0.04 | 0.72±0.04 | 1.14±0.06 | 1.16±0.06 | 1.29±0.06 | 1.28±0.07 |
| **Aripipraz.** | 0.66±0.05 | 0.64±0.04 | 0.69±0.03 | 0.70±0.05 | 1.15±0.06 | 1.16±0.06 | 1.28±0.06 | 1.27±0.07 |

### Discussion

Using a probabilistic learning paradigm with factual (where subjects were informed only about the result of the chosen outcome) and counterfactual (where subjects were also informed about the result of the unchosen outcome) feedback information conditions in healthy controls and aripiprazole-medicated and unmedicated GTS patients, we showed a specific effect of aripiprazole on counterfactual learning in the absence of any effect on factual learning.

All subjects performed significantly above chance level in the partial feedback condition, and accuracy did not differ across the groups. As previously shown, control subjects had higher accuracy in the complete feedback condition than in the partial feedback condition^22,26^. However, in GTS patients, counterfactual learning was affected by medication status: unmedicated patients behaved similarly to healthy controls, whereas aripiprazole-medicated patients did not improve their performance in the presence of counterfactual feedback. Counterfactual learning, as implemented in our task, requires at least two component processes: a “peripheral” process, which corresponds to the fact that the complete feedback condition mobilizes additional attentional resources; and a “central” process, which corresponds to the integration of hypothetical prospects (“counterfactuals”) in reinforcement learning^27^.

Was the effect of aripiprazole on counterfactual learning the hallmark of a specific impairment in counterfactual reasoning or a consequence of reduced attention? Evidence against a nonspecific “peripheral” effect is the fact that baseline performance (as measured in the partial feedback condition) did not differ in the unmedicated and the aripiprazole GTS groups. If anything, average accuracy in the partial feedback condition was numerically higher in the aripiprazole GTS group than in the unmedicated GTS group (67.5% vs. 63.0%). Further evidence against the idea of a general attentional decrease is our finding that reaction times did not differ among the groups, indicating a similar level of engagement in the task for the subjects in all groups.

The preserved ability to learn from factual outcomes during aripiprazole treatment replicates previous work by Worbe and colleagues showing that aripiprazole did not affect probabilistic reward learning in GTS patients^12^. This characteristic of aripiprazole strikingly contrasts with the robust observation that dopamine antagonists blunt reward-related learning and performance^9-11^. Reduced counterfactual learning has been reported in nicotine addiction^28^. It may therefore be argued that this learning deficit may contribute to the increased likelihood of behavioural addictions, such as pathological gambling, in aripiprazole-medicated patients^29^ as well as the observation that aripiprazole increases cigarette smoking^30^.

The reduction in counterfactual learning in the aripiprazole group may come as a surprise because early studies proposed that, given its particular mechanism of action as a partial D2 agonist, aripiprazole would preserve or even improve high-level cognitive function through improved frontal dopamine release^31,32^. However, later studies mitigated this expectation by showing no effect or even a deterioration in executive functions following aripiprazole administration^33-39^. Consistent with those findings, imaging studies failed to demonstrate an increase in frontal function and even showed reduced frontal metabolism associated with reduced performance in executive function tasks^40-42^. These contrasting findings on prefrontal dopamine might be linked to aripirazole dose; low-dose systemic aripiprazole (0.5 mg/kg) was shown to increase extracellular dopamine levels in the cortex, whereas high-dose aripiprazole (10 mg/kg) reduced dopamine levels in mice^43,44^ This might explain why we find a detrimental effect on counterfactual learning that increases significantly with aripiprazole dose in our study.

Aripiprazole has binding affinity for both dopamine and serotonin receptors^1^. Its affinity for 5-HT1A receptors in particular may provide an explanation for the apparent discrepancy between preserved factual learning and reduced counterfactual learning. Dopamine and serotonin have been associated with both automatic/reflexive and deliberative/reflective reinforcement processing, which in our task could be incorporated into factual and counterfactual learning^45-48^. In particular, it has been shown that dietary depletion of tryptophan reduces the propensity to implement cognitive strategies in a reinforcement learning task^46,49^, and aripiprazole has been shown to reduce serotonin output in the prefrontal cortex and the dorsal raphe nucleus^50^. In the light of these findings, it may be argued that aripiprazole’s detrimental effect on counterfactual learning is mediated by an alteration in serotoninergic neurotransmission. Aripiprazole would therefore have a specific pattern of effects on reinforcement learning that includes both preserved direct/factual learning due to its dopamine partial agonist property and reduced indirect/counterfactual learning due to its prefrontal serotonin disruption effect. Future studies in which specific dopaminergic and serotoninergic pharmacological agents are used are required to determine whether the effect of aripiprazole on counterfactual learning is mediated by its dopaminergic affinity or its serotonergic affinity or by a combination of the two.

Is the detrimental effect of aripiprazole on counterfactual learning specific to GTS patients or generalizable? The answer to this question is closely related to the mechanism through which aripiprazole blunts counterfactual learning. GTS has been hypothesized to be dependent on a constitutive hyperactivity of dopaminergic neurotransmission^51,52^ Positron emission tomography and post-mortem studies found evidence of increased dopamine concentration in both the striatum and the prefrontal cortex of GTS patients^53-57^. Thus, if the negative effect of aripiprazole on counterfactual learning is mediated by the dopamine system, the alterations of this neuromodulator that occur in GTS may preclude the generalizability of our result to other pathologies and to the general population. On the other hand, there is no current clear hypothesis and no replicated data implicating the serotoninergic system in GTS, with recent negative results^58,59^. Thus, if the negative effect of aripiprazole on counterfactual learning is mediated by the serotonin system, it is less likely that it is specific to GTS patients.

Previous studies suggest that hyperactive reinforcement learning may contribute significantly to GTS symptoms^60,61^. Assuming that positive and negative prediction errors are represented by phasic fluctuations in dopaminergic transmission, we previously found that reinforcement learning in unmedicated GTS subjects was consistent with a functional hyperdopaminergia, both in terms of enhanced reward^62^ and reduced punishment learning^63^. These findings were obtained when reward-outcome contingencies were deterministic and unconscious as measured by post-test assessments of cue visibility and stimulus-reward associations and using a task that has been shown to rely specifically on the ventral striatum^64^. In contrast, our current results were obtained using a probabilistic instrumental learning task that employs perfectly visible (therefore conscious) cues and recruits both the cortex (namely, the medial prefrontal cortices and the insula) and the ventral striatum^22^. In the present study, we found that unmedicated GTS patients performed equally well in the reward-seeking condition and in the punishment-avoidance condition. Taken together, our current and previous findings might suggest that in GTS excessive positive reinforcement occurs for implicit/unconscious (as opposed to explicit/conscious) learning processes; this possibility could hypothetically be supported by the dependence of these two types of processes on different neural substrates (subcortical vs. prefrontal)^65^.

A potential limitation of our study is that it was observational, meaning that we included patients who were already either unmedicated or undergoing treatment with aripiprazole.

The absence of group randomization raises the question that the effects attributed to aripiprazole may arise from pre-existing phenotypic between-group differences that determined the patients’ medication status. This is unlikely for several reasons. First, very few patients are either never or always medicated; thus, medication status fluctuates and does not seem to be a patient-specific characteristic^8^. Second, to control as much as possible for this potential confounding factor, we measured several clinical, psychological and cognitive parameters and compared them between groups. We ensured through a psychiatric interview that the included GTS subjects did not have comorbid psychiatric diagnoses. We further measured a number of dimensions that could account for learning differences in our task, including general intelligence, disease severity, and cognitive and clinical dimensions, such as impulsivity, anxiety and depression, through rating scales. None of these dimensions differed significantly between aripiprazole-treated and unmedicated GTS patients; it is therefore unlikely that they could account for the reduction in counterfactual learning found in the aripiprazole group. Two of these dimensions, depression (BDI) and level of study, differed significantly between the control group and the two GTS groups, and impulsivity differed marginally. This, of course, does not affect our main claim regarding a difference between unmedicated and aripiprazole GTS patients. However, as a formal control, we included these dimensions as cofactors in the ANOVA analysis and found that the reported effects remained significant. To summarize, while the absence of group randomization is a methodological limitation that requires us to be cautious in concluding that the difference in counterfactual learning across groups is a specific result of treatment, we controlled for a number of potential confounding factors to make this conclusion as robust as possible.

Another potential limitation of our study is its relatively small sample size. While the number of participants analysed in each group is consistent with previous publications targeting reinforcement learning in GTS patients^12-62,63^, one could wonder whether the absence of difference in accuracy between the complete feedback and partial feedback conditions in the aripiprazole group might be due to a lack of power. However, the Bayesian analysis provided evidence that the absence of a difference in accuracy between the partial and complete feedback conditions specifically in the aripiprazole GTS group did not arise as the result of a lack of power.

To conclude, we demonstrated a negative effect of aripiprazole on counterfactual learning in a population of aripiprazole-medicated GTS patients. At present, it is unclear whether this finding can be generalized to other clinical populations and, most notably, to schizophrenic patients, for whom treatment with aripiprazole is becoming increasingly popular. It is also unclear at this stage whether the observed effect is mediated by the dopaminergic system or the serotoninergic system. Further research is needed to address these questions. Nonetheless, our findings shed new light on the characterization of the cognitive profile of aripiprazole and represent a first step in understanding the neuropharmacology of counterfactual learning.

## Materials and methods

### Participants

The ethics committee of the Pitie-Salpetriere Hospital approved the study (N° Promoteur: C11-34; N° CPP: 97*19; N° ANSM: B121209-31). All participants gave written informed consent prior to participation in the study, and the study was conducted in accordance with the Declaration of Helsinki (1964).

Forty consecutive patients of the GTS Reference Center in Pitie-Salpetriere Paris were screened for inclusion and offered participation in this study. They were examined by a multidisciplinary team experienced in GTS (AH and YW). Tic severity was rated on the Yale Global Tic Severity Scale^66^. Inclusion criteria for the patients were as follows: age above 18 and confirmed diagnosis of GTS fulfilling the DSM-5 criteria. Aripiprazole-medicated GTS patients were on stable antipsychotic treatment for at least 4 weeks. Exclusion criteria included the co-occurrence of Axis I psychiatric disorders, including current major depressive disorder, current anxiety disorder, current or past psychotic disorder, substance abuse aside from nicotine, attention deficit disorder and obsessive compulsive disorders. These pathologies were screened using both subsections of the Mini International Neuropsychiatry Inventory (MINI) and a clinical evaluation performed by a trained neurologist who investigated the presence of diagnostic criteria for OCD or ADHD as detailed in the DSM-IV TR. Exclusion criteria also included any neurological or movement disorder other than tics as evaluated by a qualified neurologist before inclusion, as well as the use of any medication (including antidepressants and benzodiazepines) in the unmedicated GTS group and the presence of any co-medication in the aripiprazole GTS group.

Healthy volunteers were recruited by hospital-based advertisements. The inclusion criterion for healthy volunteers was age over 18 years. The exclusion criteria were the same as for TS patients plus a personal history of tics and any concomitant treatment except contraceptive pills for women.

### Behavioural task

The subjects performed a probabilistic instrumental learning task adapted from previous imaging studies^22,25^. The subjects were first provided with written instructions, and these were reformulated orally if needed. Each participant first performed a short training session (16 trials) aimed at familiarizing the participant with the task’s timing and responses. The participant then performed three learning sessions.

The options were abstract symbols taken from the Agathodaimon font. Each session contained eight novel options grouped into four novel fixed pairs. The pairs of options were fixed so that a given option was always presented with the same other option. Thus, within each session, pairs of options represented stable choice contexts. Within sessions, each pair of options was presented 24 times for a total of 96 trials. The four option pairs corresponded to the four conditions (reward/partial, reward/complete, punishment/partial and punishment/complete), and the four conditions were associated with different pairs of outcomes (reward contexts: winning 0.5 € versus nothing; punishment contexts: losing 0.5 € versus nothing) and the provision of a different amount of information at feedback (partial or complete). In the partial feedback contexts, only the outcome regarding the chosen option was provided, whereas in the complete feedback contexts the outcomes of both the chosen and the unchosen option were provided (Figure 1A). Within each pair, the two options were associated with the two possible outcomes with reciprocal probabilities (0.75/0.25 and 0.25/0.75). The subjects were informed that the aim of the task was to maximize their payoff and that only factual (and not counterfactual) outcomes counted. Between cues, the outcome probability was independent on a trial-by-trial basis, although it was anti-correlated on average. Thus, in a complete feedback trial, the subjects could observe the same outcome from both cues on 37.5% of trials and different outcomes from each cue on 62.5% of trials.

Pairs of options were presented in a pseudo-randomized and unpredictable manner to each subject (intermixed design). During each trial, one option was randomly presented on the left and one on the right side of a central fixation cross (Figure 1B). The side on which a given option was presented was also pseudo-randomized such that a given option was presented an equal number of times on the left and on the right of the central cross.

### Statistical analysis

From the learning task, we extracted accuracy. An “accurate” (or “correct”) choice occurs when the subject selects the most rewarding symbol in the reward-seeking context (i.e., the symbols that yielded +0.5 € in 75% of cases and 0 € in 25% of cases) or selects the less punishing symbol in the punishment-avoidance context (i.e., the symbols that yielded 0 € in 75% of cases and -0.5 € in 25%). The central hypothesis regarding the effect of aripiprazole was assessed using a repeated measures ANOVA with group (« Controls », « Unmedicated », and « Aripiprazole ») as the between-subjects factor and trial (1:24), feedback valence (reward vs. punishment avoidance) and feedback information (partial feedback vs. complete feedback) as the within-subject factors. A specific effect of aripiprazole on counterfactual learning would result in a significant group x feedback information interaction. As a control, we performed an ANCOVA analysis that included additional psychometric measures (education, BDI, BIS.11 and BIS.BAS) as covariates. As an additional control, we performed the same ANOVA on reaction times. We also performed a confirmatory ANOVA on the “counterfactual gain”, which was defined as the difference in the accuracy achieved in the complete and partial feedback contexts (a positive “counterfactual gain” indicates a beneficial effect of counterfactual feedback on performance).

To assess the directionality of the effect, we reported the average values of all dependent variables as well as the results of Student’s t-tests (two-sample t-tests were used when comparing groups, and one sample t-test was used when comparing group-values to zero). A Welch-Satterthwaite approximation was used when the variance was not equal across the groups^67^. Correlations, in particular the correlation between treatment dose and accuracy, were computed using Spearman's test. The significance level was set at 5%.

To assess evidence in favour of the null hypothesis (i.e., no difference in accuracy in the complete and partial feedback conditions), we performed a Bayesian equivalent of a t-test using Monte Carlo Markov chain analysis to compute the posterior probability of the difference in accuracy between the two feedback information conditions for each group. This analysis was conducted using the Bayesian Estimation Supersedes the t-Test (BEST) package version 0.4.0 in R statistical software with the BESTmcmc and plot.BEST functions. The region of practical equivalence (ROPE) was set at 3%, meaning that we considered a difference included in the [−1.5%, 1.5%] interval to be too small to be considered relevant. Evidence for the null hypothesis was the probability that the actual difference between feedback information conditions lies in the ROPE for each group. This probability was assessed through Monte Carlo Markov chain analysis; 100 000 simulations were run for each test.

All statistical analyses were performed using R statistical software.

